# RIPK3 coordinates RHIM domain-dependent inflammatory transcription in neurons

**DOI:** 10.1101/2024.02.29.582857

**Authors:** Sigal B. Kofman, Lan H. Chu, Joshua M. Ames, Suny Dayane Chavarria, Katrina Lichauco, Brian P. Daniels, Andrew Oberst

## Abstract

Neurons are post-mitotic, non-regenerative cells that have evolved fine-tuned immunological responses to maintain life-long cellular integrity; this includes resistance to common programmed cell death (PCD) pathways, including apoptosis and necroptosis. We have previously demonstrated a necroptosis-independent role for the key necroptotic kinase RIPK3 in host defense against neurotropic flavivirus infection. While this work showed that neuronal RIPK3 expression is essential for chemokine production and recruitment of peripheral immune cells to the infected CNS, the full RIPK3-dependent transcriptional signature, and the molecular mechanism underlying RIPK3-dependent transcription in neurons are incompletely understood. It also remains unclear what factors govern differential RIPK3 effector functions in different cell types. Here, we show that RIPK3 activation has distinct outcomes in primary cortical neurons when compared to mouse embryonic fibroblasts (MEFs) during Zika virus (ZIKV) infection or following sterile activation. We found that RIPK3 activation does not induce death in neurons; in these cells, RIPK3 is the dominant driver of antiviral gene transcription following ZIKV infection. While RIPK3 activation in MEF cells induces cell death, ablation of downstream cell death effectors unveils a RIPK3-dependent transcriptional program which largely overlaps with that observed in ZIKV-infected neurons. Using death resistant MEFs as a model to study RIPK3 signaling revealed that RIPK3 transcription relied on interactions with the RHIM domain-containing proteins RIPK1 and TRIF, effects mirrored in the RIPK3-dependent antiviral transcriptional signature observed in ZIKV-infected neurons. These findings suggest the pleotropic functions of RIPK3 are largely context dependent and that in cells that are resistant to cell death, RIPK3 acts as a mediator of inflammatory transcription.

**One Sentence Summary:** RHIM-domain containing proteins form a conserved signaling network capable of mediating inflammatory transcription and cell death.

## INTRODUCTION

Neurons of the central nervous system are post-mitotic, display limited replicative potential, and may persist for the life of the organism^1^. As such, neurons have evolved mechanisms to maintain cellular integrity both at homeostasis and in response to infection or stress. For example, neurons have been shown to implement neuron-specific, replication-independent genome repair mechanisms^2^. It has also been theorized that CNS neurons have limited regenerative capacity to preserve complex neuronal networks, as improper addition of synapses could be detrimental^3,4^. Importantly, as a component of this specialization and longevity, neurons have developed immunological responses that control early cell-intrinsic infection, thereby restricting antigen dissemination to limit the magnitude of peripheral inflammatory responses and prevent cellular damage^5,6^. Among these specialized immunological responses is reduced susceptibility to programmed cell death (PCD), including both apoptosis and necroptosis.

Necroptosis is a form of inflammatory PCD mediated by the adapter kinase RIPK3^7–9^. Once activated, RIPK3 phosphorylates the executioner of necroptosis, mixed lineage kinase domain-like protein (MLKL), leading to MLKL oligomerization and translocation to the cell membrane and subsequent membrane rupture^10,11^. RIPK3 activation occurs via formation of a cytosolic complex termed the “necrosome”; this can be initiated by a variety of upstream proteins and sensors, including TNF-family receptors acting via RIPK1, TLR3 or TLR4 acting via TIR-domain-containing adapter-inducing interferon-B (TRIF), and self- or viral nucleic acids, acting via Z-DNA binding protein 1 (ZBP1).These four proteins (RIPK3, RIPK1, TRIF and ZBP1) interact via RIP Homotypic Interaction Motifs (RHIMs), and are the only four mammalian proteins described to encode these domains^8^. The RHIM domain is essential for binding and hetero-oligomerization of RIPK1, ZBP1, and TRIF to RIPK3 to induce necroptotic cell death and transcription^12–15^. Moreover, as RIPK1, TRIF and ZBP1 are each activated by specific stimuli, it has generally been assumed these necroptotic signaling pathways are linear, with limited pathway crosstalk^7,16,17^.

While most cells undergo necroptotic cell death upon RIPK3 activation, we recently described a novel, cell death-independent role for RIPK3 in primary cortical neurons^18,19^. Our studies showed that West Nile virus (WNV)- or Zika virus (ZIKV)-infected neurons activate RIPK3 and induce RIPK3-dependent inflammatory gene transcription, including induction of anti-viral chemokines and interferon stimulated genes (ISGs), in the absence of cell death^18,19^. However, the full RIPK3-transcriptional signature and the molecular mechanism governing transcription driven by the necroptotic pathway is not well understood. Moreover, using a mouse model of flavivirus-encephalitis we also showed that RIPK3 restricted WNV and ZIKV replication independent of MLKL, and as such, the role of MLKL in neurons and the factors that determine whether RIPK3 activation leads to cell death or transcriptional outcomes (or both concurrently), remain unclear^20^.

Here, we define the RHIM-containing-protein signaling network nucleated by RIPK3 in neurons. To elucidate RIPK3 signaling we carried out RNAseq analysis on Zika virus-infected wildtype (WT) and *Ripk3^-/-^* neurons or MEF cells and found that RIPK3 is required for an anti-viral gene response only in neurons. Moreover, a similar functional inflammatory signature was also observed in sequencing analysis carried out using a RIPK3-activatable system, in which necrosome formation can be directly induced in cells, implying that RIPK3 activation is both necessary and sufficient for antiviral transcriptional responses in neurons. Surprisingly, blocking downstream cell death effectors in MEFs, which typically undergo rapid necroptosis, revealed a RIPK3-dependent transcriptional program that phenocopied RIPK3 signaling in neurons. This transcriptional response is dependent on contributions from both RIPK1 and TRIF downstream of RIPK3 oligomerization. This mechanism largely modeled RIPK3- dependent transcriptional signaling in ZIKV-infected neurons, which we found to also depend on concurrent activation of TRIF and RIPK1, in a complex initiated by ZBP1 and nucleated by RIPK3. Together, these findings suggest that RHIM-containing proteins form a signaling network defined by extensive crosstalk, that RHIM-dependent signaling is a prime driver of transcriptional responses in ZIKV-infected neurons, and that in this setting the primary output of this pathway is transcription, not death.

## RESULTS

### RIPK3-dependent inflammatory gene transcription during flavivirus infection is neuron-specific

To investigate RIPK3-dependent, cell-type specific responses to ZIKV infection, we derived primary MEFs and cortical neurons from embryonic *Ripk3^-/-^* and congenic control C57BL/6J (WT) mice. First, we infected neurons with the African lineage strain ZIKV-MR766 (Uganda, 1947) MOI 0.1 for 24 hours and harvested total RNA for RNA sequencing analysis to assess the full transcriptional response, and to define RIPK3-dependent genes, induced by ZIKV infection. Principal component analysis showed clustering among mock samples and distinct separation between infected WT and *Ripk3^-/-^* samples (Fig. S1A), suggestive of RIPK3-dependent transcriptional differences. While ZIKV-infection induced the expression of characteristic anti-viral genes in WT neurons, this signature was virtually absent in *Ripk3^-/-^* neurons (Fig. 1A, B, and C). Very few differences in gene expression were observed between mock samples (Fig. S1B). The RIPK3-dependent gene signature was heavily enriched in prototypic interferon stimulated genes (ISGs) and gene ontology (GO) analysis confirmed ZIKV-induced, RIPK3-depenendent genes in neurons had innate immune and anti-viral functions as well as being significantly over-represented in inflammatory signaling pathways by gene set enrichment analysis (GSEA) (Fig. 1D and E).

**Fig. 1:**
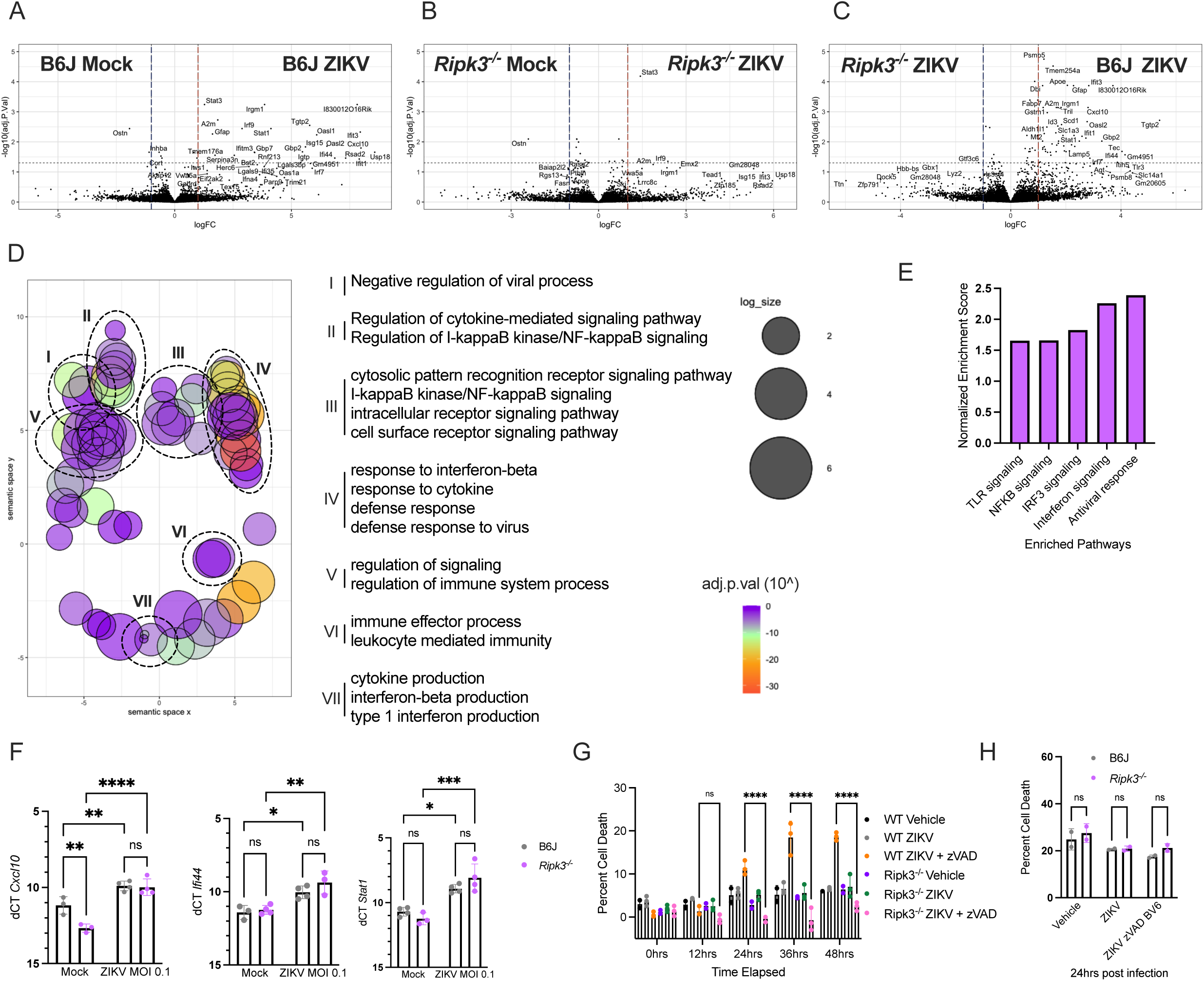
RIPK3-dependent inflammatory gene transcription during flavivirus infection is neuron-specific. **(A)** Volcano plot depicting differentially expressed genes in WT neurons 24 hours after ZIKV-MR766 (MOI 0.1) infection as compared to WT mock samples (*N*=3 biological replicates). **(B)** Volcano plot depicting differentially expressed genes in *Ripk3^-/-^* neurons 24 hours after ZIKV-MR766 (MOI 0.1) infection as compared to *Ripk3^-/-^* mock samples (*N*=3 biological replicates). **(C)** Volcano plot depicting differentially expressed genes in WT neurons 24 hours after ZIKV-MR766 (MOI 0.1) infection as compared to *Ripk3^-/-^* ZIKV samples (*N*=3 biological replicates). **(D)** GO analysis displaying proposed biological functions of genes in Figure 1D. Bubble color represents adj.p.val and log_size represents number of annotations associated with each GO ID. The x and y axes represent “semantic space”. **(E)** GSEA displaying enriched pathways in ZIKV-infected WT versus *Ripk3^-/-^* infected neurons. GSEA based on MsigDb canonical pathways collection 2 (Curated gene sets). **(F)** mRNA expression measured by qRT-PCR in primary MEFs derived from WT and *Ripk3^-/-^* mice 24 hours after ZIKV-MR766 (MOI 0.1) infection (*N*=3 biological replicates). **(G)** Percent cell death in WT and Ripk3-/- MEFs after ZIKV infection or ZIKV plus zVAD (*N*=3 biological replicates). **(H)** Percent cell death in WT and *Ripk3^-/-^* neurons after ZIKV infection or ZIKV plus zVAD and BV6 (*N*=2 Representative of 3 separate experiments). ns, not significant. *p<0.05, **p<0.01, ***p<0.001, ****p<0.0001. Error bars represent SD.

We next carried out transcriptional analysis on ZIKV-infected primary MEFs to determine whether anti-viral gene expression was RIPK3 dependent in this cell type. WT and *Ripk3^-/-^* MEFs were infected with ZIKV-MR766 MOI 0.1 for 24 hours, and RNA lysates were subsequently analyzed. As expected, ZIKV-infected MEFs significantly upregulated anti-viral genes but unlike neurons, gene expression was RIPK3-independent (Fig. 1F). Although we did not observe RIPK3-mediated induction of the genes we assessed in ZIKV-infected MEFs, we wondered whether RIPK3 was instead playing a canonical role as a cell death inducer in this setting. To address this question, we analyzed cell death kinetics in ZIKV-infected MEFs and neurons. While ZIKV alone did not induce necroptosis, de-repression of the necroptotic pathway via addition of the pan-caspase inhibitor zVAD induced cell death in a RIPK3-dependent manner in ZIKV-infected MEFs. However, ZIKV alone, in combination with zVAD, or in combination with both zVAD and the IAP inhibitor BV6 did not induce necroptosis in primary cortical neurons (Fig. 1G and H). Together, these data demonstrate distinct ZIKV-induced, RIPK3-dependent outcomes in neurons and MEFs, with RIPK3 acting as an essential transcriptional mediator but failing to induce necroptosis in neurons.

### RIPK3 activation leads to inflammatory transcription independent of cell death in neurons

We next sought to better understand the effects of RIPK3 activation in neurons. To do this, we took advantage of a system we previously developed in which RIPK3 is fused to tandem, inducibly-dimerizable FKBP^F36V^ domains to form a construct we term “RIPK3-2xFV^21,22^.” In cells expressing this chimeric protein, treatment with the cell permeable ligand “B/B homodimerizer” (hereafter referred to as “B/B”) leads to rapid RIPK3-2xFV oligomerization and activation of RIPK3. Here, we made use of mice expressing RIPK3-2xFV in the Rosa26 locus, downstream of a lox-STOP-lox cassette (Fig. 2A). By crossing these mice to animals expressing the ubiquitous Mox2-Cre, we were able to generate primary neuron or MEF cells expressing RIPK3-2xFV. As expected, B/B-induced RIPK3-2xFV activation in primary MEFs led to robust cell death. However, direct RIPK3 activation using this system did not induce cell death in primary neurons, suggesting that these cells are resistant to necroptosis (Fig. 2B). Consistent with this possibility, RIPK3 activation via the canonical stimuli TNFa, zVAD and the IAP inhibitor BV6 also led to cell death in MEFs but not neurons (Fig. 2C).

**Fig. 2:**
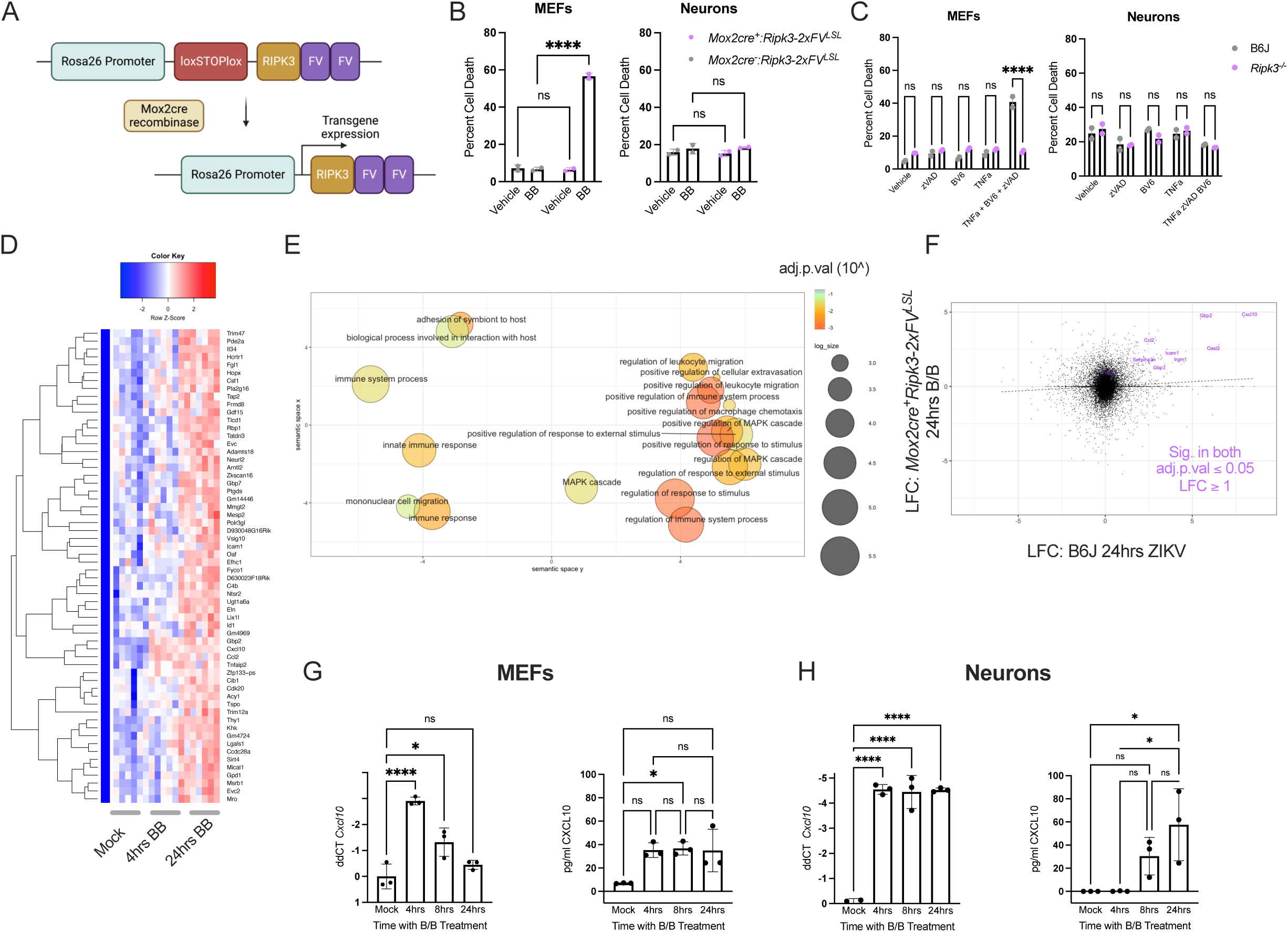
RIPK3 activation leads to inflammatory transcription independent of cell death in neurons. **(A)** Schematic for RIPK3 activatable system in mice. **(B)** Percent cell death in primary MEFs and cortical neurons derived from *Mox2cre^-^Ripk3-2xFV^LSL^* and *Mox2cre^+^Ripk3-2xFV^LSL^* mice 24 hours treatment with B/B homodimerizer (*N*=2 Representative of 3 separate experiments). **(C)** Percent cell death in primary MEFs and cortical neurons derived from WT and *Ripk3^-/-^* mice 24 hours after treatment with zVAD, BV6, TNFa, or zVAD + BV6 + TNFa (*N*=2 Representative of 2 separate experiments). **(D)** Heat map depicting row Z-scores of coordinately expressed genes in B/B treated *Mox2cre^+^Ripk3-2xFV^LSL^* neurons (*N*=6 biological replicates). **(E)** GO analysis displaying proposed biological functions of genes in Figure 2C. Bubble color represents adj.p.val and log_size represents number of annotations associated with each GO ID. The x and y axes represent “semantic space”. **(F)** Scatter plot comparing LFC values between B/B treated *Mox2cre^+^Ripk3-2xFV^LSL^* and ZIKV-infected primary neurons. Genes in purple are significantly upregulated in both conditions. **(G)** CXCL10 mRNA and protein expression measured by qRT-PCR (left) and ELISA (right), respectively, in primary *Mox2cre^+^Ripk3-2xFV^LSL^* MEFs (*N*=3 biological replicates). **(H)** CXCL10 mRNA and protein expression measured by qRT-PCR (left) and ELISA (right), respectively, in primary *Mox2cre^+^Ripk3-2xFV^LSL^* neurons (*N*=3 biological replicates). ns, not significant. *p<0.05, **p<0.01, ***p<0.001, ****p<0.0001. Error bars represent SD.

To understand whether neurons were simply resistant to necroptosis or if RIPK3 activation had alternative functions in neurons in a sterile setting, we next carried out RNA sequencing analysis on RIPK3-2xFVneurons. Sequencing analysis revealed that by 24 hours of B/B treatment, RIPK3 activation induced the expression of dozens of genes involved in leukocyte migration and innate immune functions, as characterized by GO analysis (Fig. 2D and 2E). Interestingly, while a core set of highly upregulated genes were expressed in both a RIPK3-dependent manner upon ZIKV infection and upon RIPK3-2xFV activation, other aspects of the RIPK3-2xFV gene signature were distinct from that observed upon infection (Fig. 2F). This suggested that other signals associated with infection, such as IFN priming and innate immune sensor activation, may influence the RIPK3-dependent gene expression program in neurons.

We then considered whether cell fate impacted transcriptional kinetics upon RIPK3 activation between MEFs and neurons. Both acRIPK3 MEFs and neurons expressed significant levels of CXCL10 mRNA at 4hours post B/B treatment. However, transcript levels quickly declined in MEFs but were sustained in neurons (Fig. 2G and 2H, left panels). In addition, CXCL10 protein level kinetics also differed between MEFs and neurons (Fig. 2G and 2H, right panels), with CXCL10 protein levels in cultured MEFs plateauing 4 hours after B/B treatment while levels of this cytokine in neuronal cultures were slower to rise but demonstrated continued and robust increases 24h after B/B treatment. These findings are consistent with necroptotic cell death attenuating transcription and eventually reducing translation in cultured primary MEF, but not neuronal, cells^20^. Collectively, these results suggest that RIPK3 activation drives inflammatory transcription, rather than necroptosis, in neurons.

### Exogenous MLKL is not sufficient to sensitize primary neurons to necroptosis

To understand why RIPK3 activation was not sufficient to kill neurons, we focused on neuronal expression of MLKL, as MLKL is phosphorylated by RIPK3 to drive the membrane lysis characteristic of necroptosis. Western blot analysis revealed that while MEF cells expressed MLKL as expected, MLKL protein was undetectable (3H1 antibody clone) at steady state, upon RIPK3 activation or following ZIKV infection in neurons (Fig. 3A). This observation led us to wonder whether the apparent lack of MLKL expression was sufficient to explain the failure of neurons to undergo necroptosis upon RIPK3 activation. To test this, we created adeno-associated viral particles (AAVs) encoding murine MLKL and used these reagents to exogenously express MLKL in primary neurons (or, as a control, MEF cells). AAV-mediated delivery of MLKL to wild-type neurons or MLKL-deficient MEFs was successful in inducing MLKL protein expression in both cell types as measured by Western blot, and exogenous MLKL expression in neurons was further confirmed by immunofluorescent staining (Fig. 3B, C and D). To assess whether exogenous MLKL was sufficient to induce necroptosis in MEFs, we utilized an immortalized tamoxifen-inducible RIPK3-2xFV-expressing MEF cell line, *Mlkl^-/-^Ripk3-2xFV^LSL^UbcErt2cre^+^*, which we will refer to as acR3M cells for brevity (Fig. S2A). To induce expression of RIPK3-2xFV, acR3M iMEFs were pulsed twice with 5uM of 4-hydroxytamoxifen (4OHT) and Cre-mediated excision of the stop cassette was confirmed via PCR (Fig. S2B). To confirm death resistance in acR3M iMEFs when cell death effectors are blocked or absent, we analyzed cell death kinetics in acR3M iMEFs treated with B/B homodimerizer alone or B/B homodimerizer and qVD, a pan caspase inhibitor. In these experiments, we elected to use qVD rather than zVAD because the former does not efficiently block the caspase-8/cFLIP complex, and therefore does not sensitize cells to RIPK3 activation in the manner observed with zVAD treatment^23^. As expected, acR3M iMEFs treated with B/B and qVD were completely cell death resistant, while iMEFs treated with B/B alone displayed modest levels of RIPK3-mediated apoptosis, a form of cell death observed upon RIPK3 activation when MLKL is absent (Fig. S2C)^24^. No cell death was observed upon B/B treatment of 4OHT-naïve acR3M iMEFs, as expected (Fig. S2C).

**Fig. 3:**
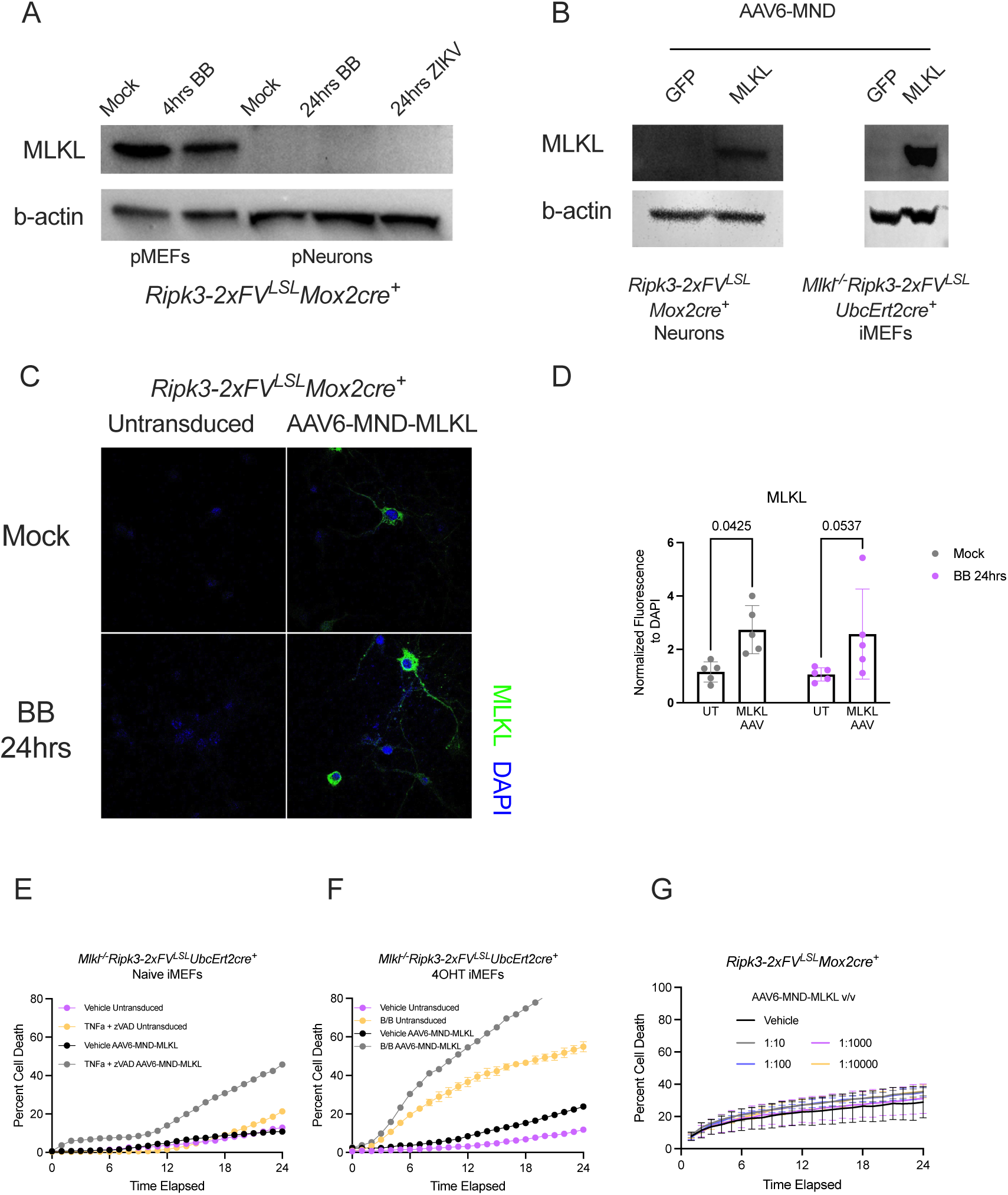
Exogenous MLKL is not sufficient to sensitize neurons to necroptotic cell death. **(A)** Detection of MLKL protein by western blot in primary *Mox2cre^+^Ripk3-2xFV^LSL^* MEFs and neurons after treatment or infection with B/B or ZIKV-MR766, respectively. b-actin used as a loading control (Representative of 2 separate experiments). **(B)** Detection of MLKL protein by western blot in acR3M iMEFs and primary *Mox2cre^+^Ripk3-2xFV^LSL^* neurons after 48 hours after AAV treatment. b-actin used as a loading control (Representative of 2 separate experiments). **(C)** Detection of MLKL protein by immunofluorescent staining and analysis in untransduced or AAV6-MND-MLKL transduced *Mox2cre^+^Ripk3-2xFV^LSL^* neurons 24 hours after B/B treatment (Representative of 2 separate experiments). **(D)** Fluorescent intensity quantification is depicted by normalizing green pixels (MLKL) to blue pixels (DAPI). (*N*=5 separate images). **(E)** Cell death kinetics of untransduced or AAV6-MND-MLKL transduced naïve acR3M MEFs treated TNFa and zVAD for 24 hours (Representative of 1 experiment). **(F)** Cell death kinetics of untransduced or AAV6-MND-MLKL transduced 4OHT acR3M MEFs treated TNFa B/B for 24 hours (Representative of 2 separate experiments). **(G)** Cell death kinetics of untransduced or AAV6-MND-MLKL transduced 4OHT *Mox2cre^+^Ripk3-2xFV^LSL^* neurons B/B for 24 hours (Representative of 2 separate experiments).

AAV-delivered exogenous MLKL was sufficient to rescue the necroptotic effects of RIPK3 activation in *Mlkl^-/-^Ripk3-2xFV^LSL^UbcErt2cre^+^*MEF cells following treatment with TNF+zVAD or upon RIPK3 activation with B/B treatment (Fig. 3E and F), confirming that AAV-mediated gene delivery results in expression of functional MLKL protein. Surprisingly however, when we tested death kinetics in AAV:MND:MLKL expressing *Ripk3-2xFV* neurons upon RIPK3 activation with B/B homodimerizer, we did not observe additional cell death over the course of 24 hours of B/B treatment in AAV-MLKL transduced acRIPK3 neurons, suggesting that exogenous MLKL expression was not sufficient to allow RIPK3 activation to induce necroptotic cell death (Fig. 3G). Together, these findings indicate that primary cortical neurons express little or no MLKL, but that even in the presence of exogenous MLKL, primary neurons resist RIPK3-dependent cell death via additional mechanisms.

### Blocking downstream cell death effectors reveals a RIPK3-dependent transcriptional program in MEFs

Since RIPK3 activation in neurons failed to induce MLKL-mediated cell death and instead drove transcriptional signaling, we wondered if ablating cell death signaling in MEFs would reveal a RIPK3-dependent transcriptional program similar to that observed in neurons. To test this, we assessed the transcriptional outputs of RIPK3 activation in acR3M cells.

To do this, we compared transcriptional and translational kinetics between acR3M cells, which succumb to RIPK3-dependent apoptosis, and the same cells co-treated with qVD to eliminate cell death responses. While death susceptible acR3M iMEFs expressed transient levels of CXCL10 mRNA similarly to acRIPK3 MEFs, addition of qVD to acR3M iMEFs phenocopied RIPK3-transcriptional kinetics seen in neurons (Fig. 4A, Left), with consistently increased levels of CXCL10 transcript up to 24 hours after RIPK3 activation. Similar results were observed upon assessment of CXCL10 protein levels (Fig. 4A, Right), together implying that cell fate rather than cell type impacts RIPK3-dependent transcription. To determine whether acR3M cells in which cell death is blocked shared a similar RIPK3-dependent transcriptional signature to that we observed in neurons, we performed RNA sequencing on acR3M iMEFs treated with B/B or B/B + qVD for 4 or 24 hours. RIPK3 activation in the absence of cell death induced robust gene expression at 4 hours which was sustained and amplified by 24 hours in iMEFs (Fig. 4B). The effect of cell death inhibition on RIPK3 transcriptional output was further emphasized when comparing gene expression magnitude between B/B alone and B/B + qVD samples 24 hours after treatment. Here, acR3M iMEFs treated with B/B + qVD upregulated significantly more genes than cells treated with B/B alone (Fig. 4C). GO analysis also ascribed R3M regulated genes with anti-viral and innate immune functions (Fig. 4D). Lastly, comparison of genes induced in a RIPK3-dependent manner upon ZIKV infection in neurons and upon RIPK3-2xFV activation in iMEFs showed considerable overlap in gene expression, though overall LFC was generally lower in MEF cells, as well as overlap in enriched pathways assessed by GSEA (Fig. 4E and 4F). These findings suggest that elimination of downstream cell death effectors unveils a RIPK3-dependent transcriptional program in MEFs that shares many features with the RIPK3-dependent program induced by ZIKV infection of neurons.

**Fig. 4:**
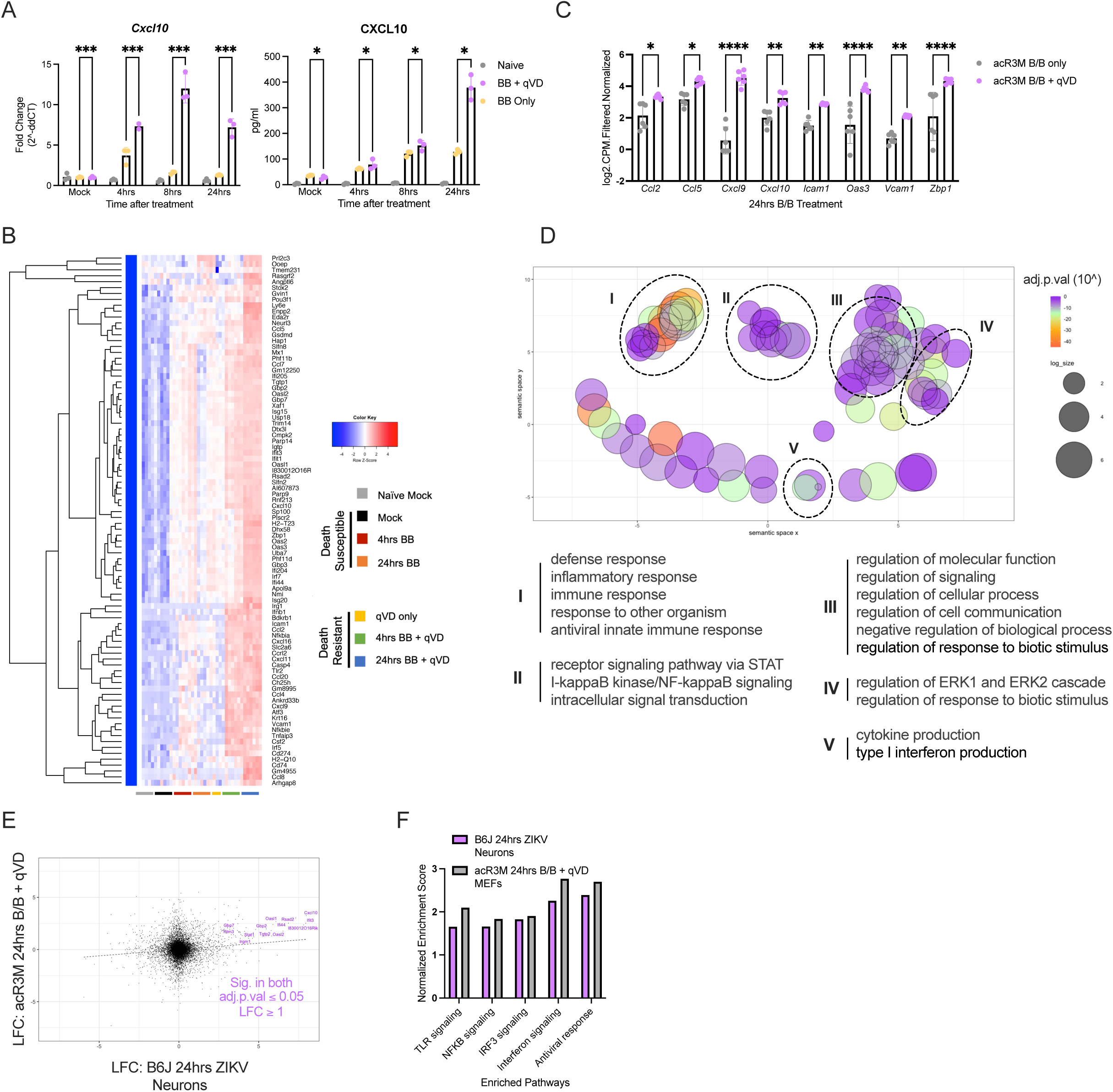
Blocking downstream cell death effectors reveals a RIPK3-dependent transcriptional program in MEFs. **(A)** CXCL10 mRNA and protein expression measured by qRT-PCR in acR3M iMEFs treated with B/B or B/B and qVD. Fold change calculated based on each condition’s mock values (Left). CXCL10 mRNA and protein expression measured by ELISA in acR3M iMEFs treated with B/B or B/B and qVD. Fold change calculated based on each condition’s mock values (Right) (*N*=3 biological replicates). **(B)** Heat map depicting row Z-scores of coordinately expressed genes in acR3M iMEFs 4 or 24 hours after B/B or B/B and qVD treatment (*N*=6 biological replicates). **(C)** Normalized and filtered Log2 CPM gene count comparison between B/B alone or B/B and qVD treated acR3M MEFs (*N*=6 biological replicates). **(D)** GO analysis displaying proposed biological functions of genes in Figure 4D. Bubble color represents adj.p.val and log_size represents number of annotations associated with each GO ID. The x and y axes represent “semantic space”. **(E)** Scatter plot comparing LFC values between B/B and qVD treated acR3M MEFs and ZIKV-infected primary neurons. Genes in purple are significantly upregulated in both conditions. **(F)** GSEA displaying overlap in enriched pathways between acR3M iMEFs treated with B/B and qVD and ZIKV-infected neurons. GSEA based on MsigDb canonical pathways collection 2 (Curated gene sets). *p<0.05, **p<0.01, ***p<0.001, ****p<0.0001. Error bars represent SD.

### Cytosolic RIPK3 activation drives RIPK1- and TRIF-transcriptional programs when cell death is blocked

As cell death resistant iMEFs phenocopied RIPK3-dependent transcription seen in neurons, we reasoned these cells could be used as a model to understand the mechanism underlying the RIPK3 transcriptional program. TRIF and RIPK1 are well established signaling mediators that drive death receptor-mediated NF-κB activation or TLR3/4- mediated IRF3 signaling, respectively^9,25,26^. Moreover, RIPK1 and TRIF are RHIM domain-containing proteins that can directly bind to RIPK3. We therefore hypothesized that RIPK3 activation could directly recruit RIPK1 and/or TRIF to drive transcriptional responses. To test this idea, we transiently suppressed expression of *Ripk1*, *Ticam1* (the gene encoding TRIF), as well as *Zbp1* (to include all RHIM domain containing proteins in our study) via siRNA in acR3M iMEFs, then performed RNA sequencing analysis on RNA samples after 4 hours of B/B and qVD treatment. We chose to carry out this experiment using transient siRNA transfection rather than CRISPR or germline knockout because we have observed that sustained ablation of RHIM proteins can alter basal transcriptional state and cellular behavior. Non-targeting siRNA pool (si*Scr*) was used as a transfection reagent control and knockdown efficacy was determined by qRT-PCR (Fig. S3A and B). Cells receiving si*Zbp1* displayed largely unperturbed RIPK3 transcriptional signaling; this was not unexpected, since ZBP1 is expressed at low basal levels and is primarily interferon-inducible (Fig. 5B)^27^. Conversely, knockdown of *Ripk1* or *Ticam1* significantly reduced expression of RIPK3 transcripts (Fig. 5C, D). These findings suggested that both RIPK1 and TRIF contribute to transcriptional activation upon RIPK3 oligomerization. To test this idea directly, we performed RNA sequencing on acR3M cells in which both *Ticam1* and *Ripk1* were knocked down, following 4 hours of B/B and qVD treatment. Strikingly, we found that RIPK3-dependent transcription was completely ablated upon knockdown of *Ticam1* and *Ripk1*, indicating that both TRIF and RIPK1 are necessary to drive RIPK3-dependent transcription (Fig. 5E and F). Gene set enrichment analysis (GSEA) revealed that RIPK3 activation upon siRNA-mediated knockdown of TRIF led to an enrichment of genes activated by NF-kB, while siRNA-mediated knockdown of RIPK1 caused enrichment of STAT5-associated genes as well as subsets of NF-kB-dependent genes (Fig. 5G). Together, these findings suggest that the RIPK3- dependent transcriptional signature is defined by TRIF and RIPK1-mediated transcription, and that each of these mediators drives a distinct component of the RIPK3-dependent transcriptional program.

**Fig. 5:**
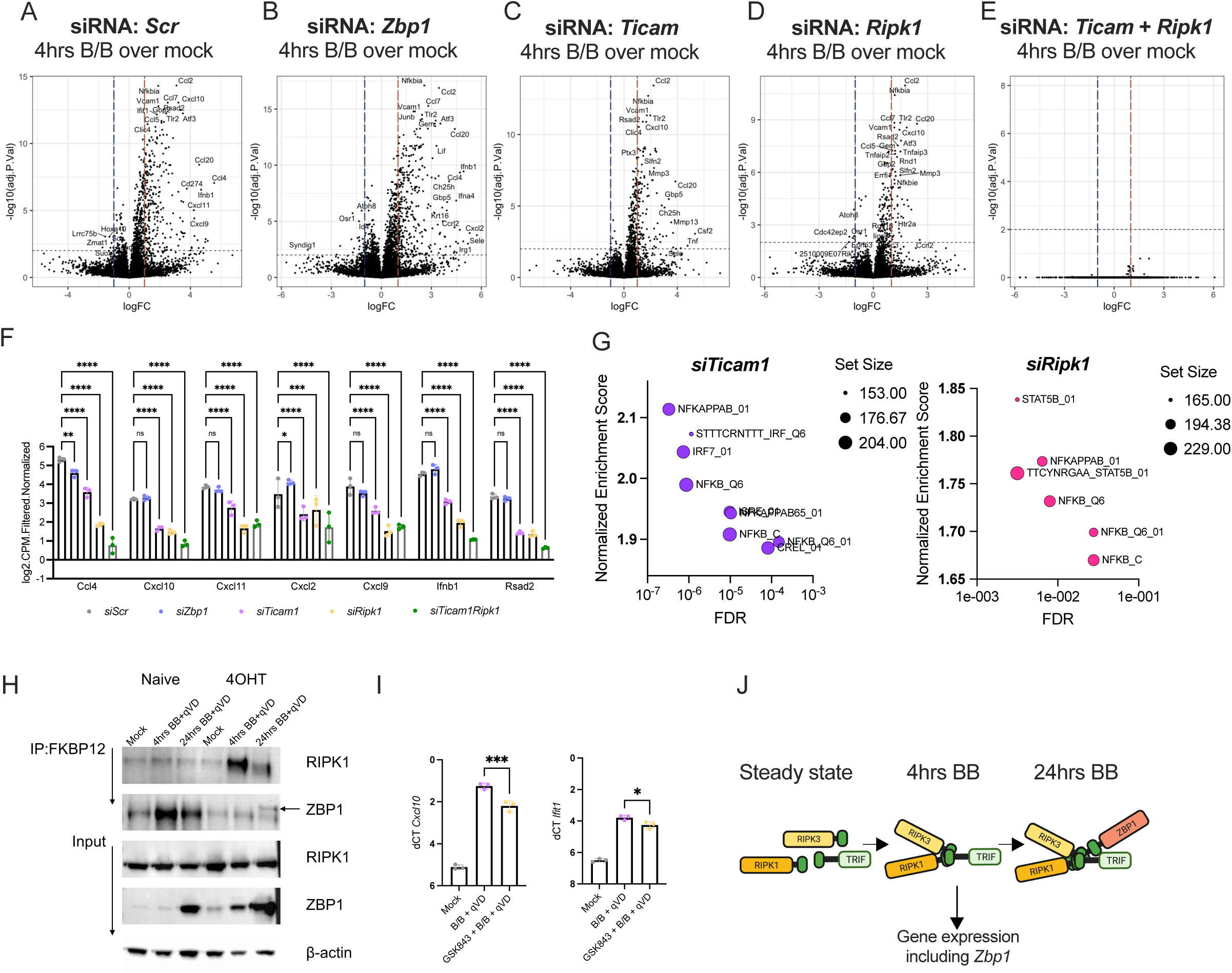
Cytosolic RIPK3 activation drives RIPK1- and TRIF-transcriptional programs when cell death is blocked. **(A-E)** Volcano plot depicting differentially expressed genes in (A) *siScr* (B) *siZbp1* (C) *siTicam1* (D) *siRipk1* and (E) *siTicam1 and Ripk1* acR3M iMEFs 4 hours after B/B and qVD treatment as compared to each condition’s respective mock samples (*N*=3 biological replicates). **(F)** Normalized and filtered Log2 CPM gene count comparison between siRNA treated acR3M iMEFs 4 hours after B/B and qVD treatment as compared to each condition’s respective mock samples (*N*=3 biological replicates). **(G)** GSEA displaying enriched pathways in si*Ticam1* (Left) and si*Ripk1* (Right) acR3M iMEFs 4 hours after treatment with B/B and qVD. GSEA based on MsigDb canonical pathways collection 3 (Curated gene sets). **(H)** Detection of RIPK1 and ZBP1 protein via western blot before and after FKBP12 immunoprecipitation (Representative image of 3 separate experiments). **(I)** mRNA expression measured by qRT-PCR in MEFs 4 hours after treatment with B/B and qVD or B/B, qVD and GSK843 (*N*=3 biological replicates). **(J)** Schematic depicting RHIM-protein complex formation after cytosolic RIPK3 activation. *p<0.05, **p<0.01, ***p<0.001, ****p<0.0001. Error bars represent SD.

To corroborate our RNA sequencing results, we carried out immunoprecipitation analysis to confirm that RIPK3 complexed with RIPK1 and TRIF. AcR3M iMEFs were treated with B/B and qVD for 4 and 24 hours, after which protein lysates were harvested and FKBP12 protein (the FV domain appended to RIPK3) was immunoprecipitated. As expected, RIPK1 bound to RIPK3 at 4 and 24hrs (Fig. 5H). Interestingly, we observed robust induction of ZBP1 expression in MEFs by 24 hours of RIPK3 activation (Fig. 4E), suggesting a positive feedback loop in which RIPK3 activation drives ZBP1 upregulation. Consistent with this observation, we found that ZBP1 interacted with RIPK3 only at later timepoints after RIPK3 oligomerization. Despite extensive testing, we were unable to identify a valid antibody for murine TRIF and were therefore unable to probe for TRIF. However, our data suggest that a cytosolic TRIF-RIPK3 complex is likely responsible for the striking transcriptional phenotype observed when *Ticam1* (TRIF) was knocked down (Fig. 5E).

Since RIPK3 has both a kinase domain and RHIM domain, we next sought to determine whether RIPK3-RIPK1 and RIPK3-TRIF signaling was driven by kinase activity or RHIM domain heterooligomerization. We thus blocked RIPK3 kinase activity with the RIPK3 inhibitor GSK843 and analyzed gene expression after 4 hours of B/B and qVD treatment in acR3M iMEFs. Inhibition of RIPK3-kinase activity moderately but significantly reduced gene expression suggesting that RIPK3 interaction with TRIF and RIPK1 is primarily RHIM domain-dependent (Fig. 5I). Together these data indicate cytosolic RIPK3 initiates transcription via TRIF and RIPK1 through RHIM domain heterooligomerization to induce the production of inflammatory proteins (Fig. 5J).

### RHIM domain proteins are required for inflammatory gene transcription in ZIKV-infected neurons

We next sought to assess the role of the RHIM domain-containing proteins in the RIPK3-dependent transcriptional program we previously observed in ZIKV-infected neurons (Fig. 1). To do this, we carried out siRNA-mediated knock-down of *Ripk3*, *Zbp1*, *Ripk1* and *Ticam1* in WT primary cortical neurons which were subsequently infected with ZIKV MR766 MOI 0.1 for 24 hours. Non-targeting siRNA pool (si*Scr*) was used as a transfection reagent control and comparisons between all mock samples was carried out to assure transcriptional baselines were comparable amongst different siRNA conditions (Fig. S4A, B, and C). RNA sequencing analysis was then carried out on total RNA from neuronal samples. As previously observed in RIPK3 knockout neurons, ZIKV-induced transcription was primarily RIPK3 dependent in siRNA treated samples (Fig. 6A and B). Neurons treated with si*Zbp1* closely phenocopied the loss of gene upregulation observed in si*Ripk3*-treated cells, supporting our previously published findings that ZIKV is sensed by ZBP1 during infection (Fig. 6A and B). Knocking down Ticam1 also significantly reduced ZIKV-induced gene expression in neurons (Fig. 6A and B).

**Fig. 6:**
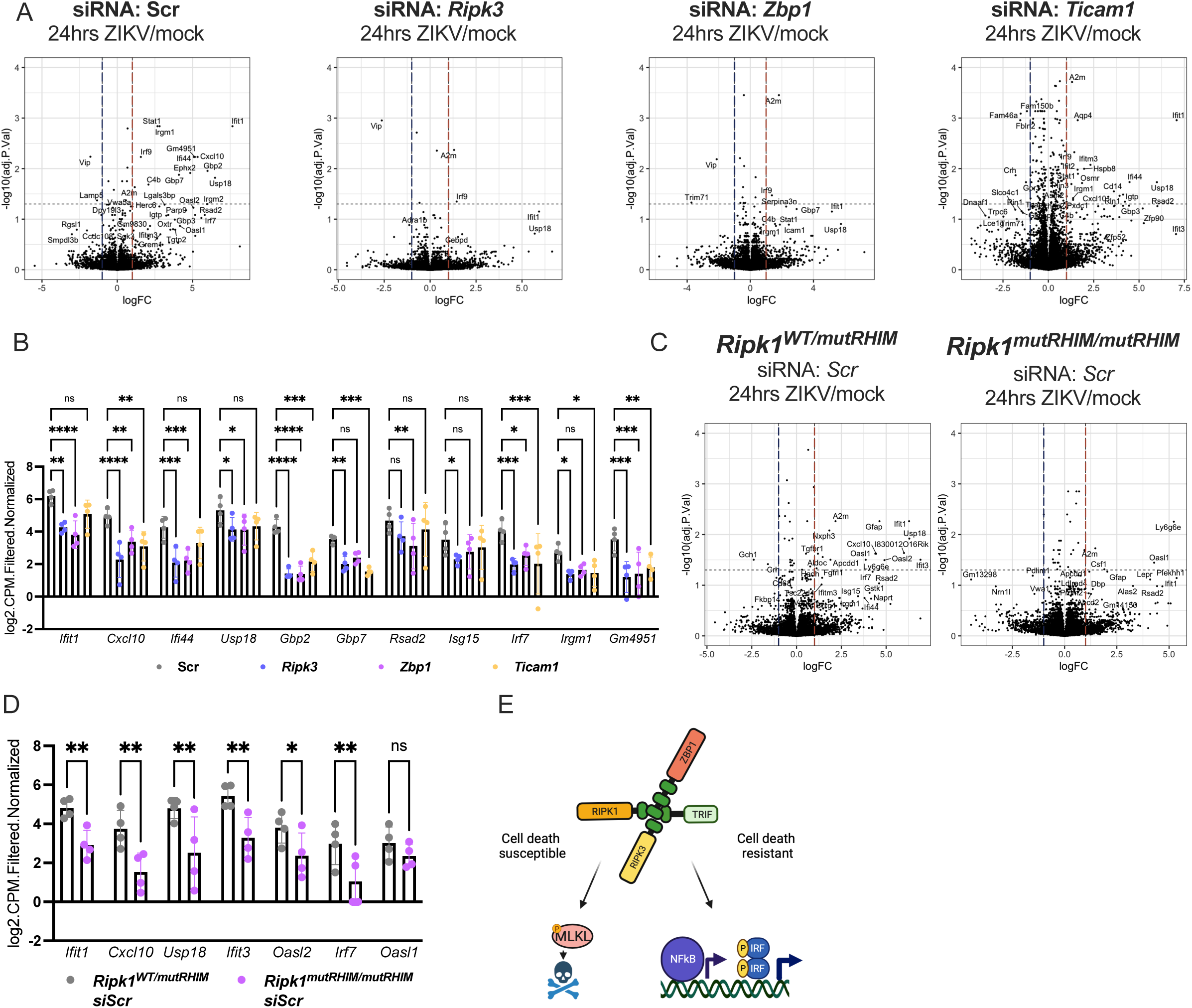
RHIM-domain proteins are required for inflammatory gene transcription in ZIKV-infected neurons. **(A)** Volcano plot depicting differentially expressed genes in *siScr*, *siRipk3*, *siZbp1* and *siTicam1* treated, ZIKV-infected neurons as compared to each condition’s respective mock control after 24 hours (*N*=4 biological replicates). **(B)** Normalized and filtered Log2 CPM gene count comparison between siRNA treated, ZIKV-infected neurons as compared to each condition’s respective mock samples (*N*=4 biological replicates). **(C)** Volcano plot depicting differentially expressed genes in *siScr* treated *Ripk1^WT/mutRHIM^* or *Ripk1^mutRHIM/mutRHIM^*ZIKV-infected neurons as compared to each respective mock control after 24 hours (*N*=4 biological replicates). **(D)** Normalized and filtered Log2 CPM gene count comparison between *siScr* treated, ZIKV-infected *Ripk1^WT/mutRHIM^* and *Ripk1^mutRHIM/mutRHIM^* neurons as compared to each condition’s respective mock samples (*N*=3 biological replicates). **(E)** Schematic depicting RHIM-protein signaling complex and possible outcomes. ns, not significant. *p<0.05, **p<0.01, ***p<0.001, ****p<0.0001. Error bars represent SD.

Unlike MEFs, we observed that neurons sustained significant changes to basal transcription upon knockdown of RIPK1, changes which were further magnified upon ZIKV-infection (Fig. S4D and E). These results were not completely unexpected, as RIPK1 has a complex and multifaceted role as both an enhancer and a suppressor of transcriptional and cell death responses and is the only RHIM-containing protein whose germline ablation is embryonically lethal^26^. To circumvent this issue and isolate the RHIM-dependent effects of RIPK1 from its other functions, we generated neurons from mice that were either heterozygous (*Ripk1^WT/mutRHIM^*) or homozygous (*Ripk1^mutRHIM/mutRHIM^*) for an inactivating mutation in the RIPK1 RHIM domain^28^. Because siRNA transfection modestly suppresses global gene expression, we treated these neurons with a scramble siRNA to allow direct comparison to the conditions described in (Fig. 6A). As expected, upon ZIKV-infection, *Ripk1^mutRHIM/mutRHIM^* neurons displayed a significant reduction in gene expression compared to infected *Ripk1^WT/mutRHIM^*(Fig. 6C and D). Collectively these results identify ZBP1 as the predominant initiator of innate immune transcriptional signaling in ZIKV-infected neurons and suggest that ZBP1 activation by ZIKV can drive transcriptional signaling via a complex nucleated by RIPK3 and containing both TRIF and RIPK1.

## DISCUSSION

Since its discovery, necroptotic cell death has been associated with inflammation. However, the source and consequences of the inflammatory response to necroptosis has remained incompletely understood. We and others have highlighted the connection between activation of the “necroptotic” pathway and concurrent inflammatory transcription programs and suggested that the latter is a primary driver of the immune response induced by activation of this pathway^14,29–33^. Our previous work has also indicated that in some cell types-notably neurons-the cell death activity of the “necroptotic” pathway is curtailed, while its transcriptional signaling remains intact^19^. However, the mechanism and breadth of this transcriptional signaling has not previously been defined. Here, we show that RIPK3 activation has distinct outcomes in primary MEFs and cortical neurons during infection and sterile inflammation. RIPK3 activation primarily induces canonical necroptotic cell death in MEFs but fails to do so in neurons. Rather, RIPK3 is an essential driver of inflammatory transcription in neurons, with RNA sequencing analysis uncovering genes for which RIPK3 is necessary as well as those for which its activation is sufficient to drive expression upon infection or sterile RIPK3 oligomerization.

Our group has previously shown that ZBP1 contributes to viral restriction following ZIKV infection in neurons, but the near absence of a transcriptional response upon knockdown of ZBP1 or RIPK3-a finding suggesting that ZBP1 is the predominant innate immune sensor activated in this setting-was unexpected^18^. Moreover, the discrepancy in transcript LFC in si*Trif* treated neurons compared to si*Zbp1* and si*Ripk3* implies that ZBP1 is the apical sensor in ZIKV-infected neurons, leading to activation of RIPK3 which then recruits RIPK1 and TRIF via RHIM-RHIM interactions, instead of signal initiation from other nucleic acid sensors. This finding further supports our hypothesis that transcriptional responses initiated by RHIM domain-containing proteins is a dominant response to ZIKV infection in neurons.

Our sequencing analysis revealed RIPK3 to be a crucial anti-viral transcriptional mediator in neurons during ZIKV-infection and sterile inflammation; notably however, we did not observe a central role for RIPK3-dependent transcription upon ZIKV infection of MEF cells, which likely rely on RIG-I-like or Toll-like receptors to drive the inflammatory response to ZIKV^34–36^. The evolutionary pressures that have led to the centrality of the ZBP1-RIPK3 pathway in sensing viral infection in neurons remain unclear, but the long co-evolution of neurotropic herpesviruses with the mammalian immune system seems a likely driver. Indeed, murine cytomegalovirus (MCMV) and herpes simplex virus 1 (HSV-1) contain viral RHIM-domains that inhibit RIPK-mediated cell death^37,38^. The degree to which these viral effectors also alter RHIM-dependent inflammation is not well understood. Interestingly, recent studies suggest that HCMV blocks necroptotic signaling downstream of necrosome formation, an effect we would predict to leave the inflammatory pathways driven by RHIM-RHIM interactions intact^39,40^. As many herpesviruses take advantage of host cell inflammatory transcription for their own propagation, blocking cell death while leaving inflammation intact could represent a beneficial adaptation by HCMV^41^.

Our finding that MLKL protein cannot be detected in primary cortical neurons but is detected in MEFs can partially explain the differential effector functions of RIPK3 activation in each cell type. However, re-expressing MLKL via AAV gene delivery was not sufficient to render neurons sensitive to RIPK3-initiated necroptosis, implying that neurons resist necroptosis via additional means. Recent studies have shed light on the mechanisms that regulate MLKL trafficking and membrane rupture, including ESCRT-III which can delay necroptotic cell death by facilitating pMLKL exocytosis and promoting membrane repair^42^. It has also been shown that pMLKL requires phosphorylated inositol phosphate (IP) rich membrane regions to translocate to the plasma membrane and induce cell lysis^43^. It is therefore possible that ESCRT-III or differential expression of the IP kinases, IPMK and ITPK1, contribute to MLKL-dependent cell death resistance in our neuronal culture system and that neurons maintain stricter control over these MLKL checkpoints, although we have not directly investigated this hypothesis in our studies. It is important to note that several studies have pointed to a deleterious role of necroptosis and MLKL in neurodegenerative disease (NDD) in mice and humans, while others have suggested that necroptotic signaling is dispensable for the development of NDD^44–49^. Interestingly, humans with an MLKL-deficiency unexpectedly develop progressive NDD, suggesting that the upregulation and activity of MLKL in neurons may therefore be context and species dependent^50,51^.

While we were unable to induce necroptotic cell death in neurons, blocking cell death effectors in MEFs unveiled a RIPK3-dependent transcriptional program that was largely absent in death-susceptible MEFs. Consistent with our neuronal RNAseq data, RIPK3 induced genes involved in anti-viral and immune defense in MEFs. These findings imply a key factor in the magnitude and duration of RIPK3 transcription is the presence or absence of downstream effector proteins, like MLKL and caspase-8, and support the idea that necroptotic cell death may curtail, rather than contribute to, inflammation^52^. This concept is also supported by work showing that cells can continue to translate protein after “death” for a limited window, suggesting that RIPK3-mediated cytokine production is inextricably tied to its role as a cell death inducer^20^.

Using our activatable RIPK3 system in MEFs, we found that cytosolic oligomerization of RIPK3 can activate both TRIF and RIPK1. This revealed that RHIM-domain proteins can aggregate independently of upstream stimuli and induce inflammatory transcription as an inherent function of this complex. We posit that RIPK1 and TRIF control different inflammatory sub-programs downstream of RHIM complex formation, given the well-described roles of RIPK1 and TRIF in promoting NF-kB and IRF-3-mediated transcription, respectively. While we observed some overlap between NF-kB and IRF gene signatures, initiating multiple defense programs is a beneficial strategy for pathogen clearance. Consistent with our results in MEFs, we also found that ZIKV-induced anti-viral transcription was RHIM-domain dependent in neurons, positioning RHIM-domain signaling as a conserved network capable of inducing inflammatory transcription when death effectors are absent.

Previous studies have demonstrated the capacity of RHIM-domain protein complexes to induce cell death and inflammation. Specifically, ZBP1 was recently shown to be required for TRIF-dependent pyroptosis and inflammatory transcription by translocating RIPKl to the “TRIFosome” during infection and sterile inflammation^53,54^. These findings, along with our data indicating that all four RHIM-domain containing proteins complex together, suggest that RHIM-domain proteins function as a network that can integrate multiple inputs (via ZBP1, TRIF or RIPK1) and mediate several effector functions including IRF- and NF-kB-dependent transcription and various cell death modalities depending on the landscape of protein constituents and downstream effectors (Fig. 6E). This concept underscores the pleiotropic and redundant responses to infection that have been driven by host-pathogen evolutionary competition and helps to explain why disruption of RHIM signaling is an evasion strategy favored by both bacteria and viruses.

We also suggest that the propensity to die by necroptosis and the regenerative potential of a given cell type may vary inversely with one another. It has been previously shown that liver-specific *Caspase-8* deletion is well tolerated in mice, but that in livers lacking caspase-8 recovery after partial hepatectomy is accompanied by supraphysiological levels of cellular proliferation as well as chronic inflammation^55^. These results suggest that fully differentiated hepatocytes do not activate the necroptotic pathway upon Caspase-8 deletion, but regenerating cells do. CNS neurons are largely non-regenerative cells whose widespread death would be detrimental to the host. Our studies compare two distinct cell types, but our understanding of the necroptotic potential of other fully differentiated cell types is limited. The inverse relationship between susceptibility to PCD and regeneration capacity could be further tested by studying the effects of RIPK3 activation in neuronal progenitor cells (NPCs) and glial progenitor cells (GPCs) compared to fully differentiated neurons or glial cells such as astrocytes or oligodendrocytes, which could display a more subtle relationship between cell death and transcription.

## MATERIALS AND METHODS

### Experimental Model and Subject Details

#### Mice

C57BL/6J, *Ripk3^-/-^*, *Ripk1^mutRHIM/mutRHIM^*, and RIPK3-2xFV^fl/fl^Mox2-cre^+^ in this study were bred and housed under specific-pathogen free conditions at the University of Washington. All mouse strains were congenic to C57BL/6J background; in all cases wild-type controls of appropriate sub-strain were used. Genotyping for RIPK3-2xFV transgene and Mox2-cre expression was carried out as previously described^19^.

#### Cell culture and infections

Primary cultures of cerebral cortical neurons and embryonic fibroblasts (MEFs) were generated and maintained using E15.5 embryos, as described^19^. Primary cortical neuron cultures were infected with ZIKV MR766 MOI 0.1. Immortalized *Ubc-ERT2cre^+^:Ripk3-2xFV^fl/fl^:Mlkl^-/-^* MEFs were generated in our lab and maintained in DMEM supplemented with 10% FBS, sodium pyruvate and HEPES. To induce deletion of stop cassette and expression of RIPK3-2xFV transgene, iMEFs were pulsed twice with 5uM of 4-hydroxytamoxifen (4OHT) over 5 days. Confirmation of stop cassette deletion was accomplished by PCR amplification of the Rosa26 locus (Supplemental Figure 2A).

#### Viruses and virological assays

ZIKV strain MR766 was provided by the World Reference Center for Emerging Viruses and Arborviruses (WRCEVA). Viral stocks were generated by infecting Vero cells (MOI 0.01) and harvesting supernatants at 72hrs.

### Method Details

#### Cell death assays

Cell death was measured and analyzed using an Incucyte imaging system. Briefly, cell death was determined by the intracellular presence of the cell-impermeable DNA-intercalating dye SYTOX Green in cultured cells. Cell death was quantified as a percentage of SYTOX Green positive cells with respect to total cell number (SYTO Green positive cells).

#### siRNA Transfection and gene knockdown

Genetic knockdown was achieved using DharmaFECT transfection reagents and siRNA transfection protocol. Primary cortical neurons and iMEFs were treated with 25nM of SMARTpool siRNA cocktails and DhamaFECT reagent for 48hrs. After 48hrs, cells were replaced in fresh media and subsequent experiments were carried out. Efficiency of genetic knockdown was assessed via qRT-PCR.

#### Adeno-associated virus (AAV) production

Briefly, HEK 293T cells were transfected with adenoviral helper HgT1, serotype pRepCap6, vector plasmid of interest and PEI transfection reagent for 48hrs. After 48hrs, cells were collected and resuspended in cell lysis buffer and lysed in three freeze-thaw cycles using LN2 and 37°C water bath. Cell lysates were treated with Thermo Universal Nuclease at 100U/ml for 30 minutes at 37°C. Cell lysates were subsequently spun down and virus was isolated using an iodixanol gradient (67,000 RPM for 1hr at 18°C). Vector stocks were aliquoted and stored at −80°C.

#### Western blot and immunoprecipitation

Primary cortical neurons, pMEFs, and iMEFs were washed with ice cold 1XPBS and lysed with ice cold 1XRIPA lysis buffer (10mM Tris-HCL pH 8.0, 1mM EGTA, 2mM MgCl2, 0.5% Triton X-100, 0.1% NaDOC, 0.5% SDS, 90mM NaCl) supplemented with Roche cOmplete Mini protease inhibitor cocktail and Roche PhosSTOP, for 30 minutes on ice. Samples were then boiled in 4XLaemli buffer (8% SDS, 240mM Tris pH 6.8, 40% glycerol 0.04% Bromophenol Blue) and freshly added 10% BME at 95C for 15 minutes. 30 micrograms of protein were run on Bolt 4-12% Bis-Tris Plus gels with Bolt™ MOPS SDS running buffer for 45 minutes at 180V/90mAMPs and transferred onto PVDF membrane at 400mAmps for approximately 1hr in Bolt transfer buffer. PVDF membranes were blocked with 5% BSA or dry milk in TBS+1%Tween-20 (TBST) for 1hr at room temperature. Membranes were stained overnight at 4C with primary antibody in 5% BSA or dry milk in TBST. Membranes were then washed three times with TBST and stained with HRP-conjugated secondary antibody for 1hr at room temperature. Membranes were treated and developed with Pierce ECL Western Blotting Substrate or SuperSignal West Femto Maximum Sensitivity Substrate.

For immunoprecipitations (IP), FKBP12 antibody was conjugated to Protein G DynaBeads at a ratio of 1ug of antibody to 6ul of Protein G DynaBeads in IP lysis buffer (25mM Tris-HCL pH 8.0, 150mM NaCl, 1% v/v Triton X-100, 10% v/v glycerol, 0.01% w/v SDS) overnight at 4C. iMEFs were washed in ice cold 1XPBS and lysed with ice cold IP lysis buffer for 30 minutes on ice. 300ug of protein lysate was treated with conjugated FKBP12 (1ug antibody/50ug protein) and incubated on tabletop shaker overnight at 4C. Concurrently, 30ug of “pre-IP” sample were boiled in 4XLaemli buffer and 10% BME and stored in −80C until all samples were run. IP samples were then washed three times with IP lysis buffer and subsequently boiled in 4XLaemli buffer and 10% BME. All samples were run, transferred, and developed as previously described using the Bolt western blot reagents.

#### ELISA for protein analysis

CXCL10 expression in cell culture supernatants was measured using Invitrogen IP-10 (CXCL10) mouse ELISA kit and carried out per manufacturers instructions. ELISA plates were read on BioTek Synergy HT plate reader using Gen5 software.

#### Immunofluorescence staining and analysis

After treatment, cells were washed with ice-cold PBS and fixed with 100% methanol for 30 minutes. Cells were blocked and permeabilized for one hour in ice-cold TBS (20nM Tris, 150nM NaCl, pH 7.6) + 0.05% v/v Triton-X100 and 10% donkey serum and subsequently incubated with rat-MLKL (Millipore MABC604 clone 3H1 1:1000) overnight. Cells were incubated in secondary antibody donkey anti-rat 488 (ThermoFischer Scientific A21208) for three hours. Nuclei were visualized using Vectashield mounting media with DAPI. Images were analyzed with FIJI software.

#### RNA isolation and gene expression analysis

Total RNA from cell cultures was isolated using a Macherey-Nagel NucleoSpin RNA kit. cDNA was synthesized using Invitrogen SuperScript III Reverse Transcriptase and gene expression was analyzed via quantitative reverse transcriptase (qRT-PCR) using Sybr Green reagents and a ViiA 7 Real-Time PCR system. Delta cycle threshold (CT) values were quantified by normalizing to the CT value of housekeeping gene GAPDH (CT_target_-CT_GAPDH_). Data was further normalized to baseline controls of respective genotypes (delta CT_target(genotype x)_ - delta CT_control (genotype x))._

#### RNA-sequencing analysis

Primary cortical neuron and iMEF cultures were sequenced at the Benaroya Research Institute Genomics Core Laboratory. Quality of RNA was determined on a TapeStation, and libraries were prepared using NexteraXT library preparation kit. Samples were run on an Illumina NextSeq2000, to generate 59 base-pair, paired-end reads with a depth of approximately 5 million reads per sample. Sequencing data were analyzed using the DIY.transcriptomics (diytranscriptomics.com) pipeline using R and Rstudio programming. Briefly, raw reads were mapped to the Genome Reference Consortium Mus musculus 39 using Kallisto. Quality of reads was assessed with FASTqc and Multiqc. Transcripts were annotated using Ensembl database Mus musculus version 79. Samples were filtered to exclude genes with counts per million = 0 in 6 or more samples (iMEFs) or 4 or more samples (neurons), and log2 counts per million were subsequently normalized using EdgeR. The Voom function in the LIMMA package was used to variant stabilize our data. LIMMA was used to apply a linear model to our data and Bayesian statistics were used to determine significantly up- or down-regulated genes by a log fold change of 1 and false-discovery-rate of 0.01. The open-source gene ontology analysis platform, “REVIGO: reduce + visualize gene ontology” source was used to create semantic space plots (Supek F, Bosnjak M, Skunca N, Smuc T. “REVIGO summarizes and visualizes long lists of Gene Ontology terms” PLoS ONE 2011. doi:10.1371/journal.pone.0021800). Enriched pathways identified via Gene Set Enrichment Analysis (GSEA) were abbreviated for clarity. The key is as follows: WP_TOLLLIKE_RECEPTOR_SIGNALING_PATHWAY (TLR signaling), HINATA_NFKB_TARGETS_KERATINOCYTE_UP (NFKB signaling), GRANDVAUX_IRF3_TARGETS_UP (IRF3 Signaling), BOSCO_INTEFERON_INDUCED_ANTIVIRAL_MODULE (Antiviral Signaling), REACTOME_INTERFERON_SIGNALING (Interferon Signaling).

### Statistical Analysis

Statistical analysis was completed using GraphPad Prism 10. Experimental data was compared using parametric testing, including 2-tailed student’s t test, l or 2-way ANOVA with appropriate multiple comparisons test. All data points represent biological replicates unless noted otherwise.

## Supplementary Material

Figure S1. RNA sequencing analysis of ZIKV-infected WT and *Ripk3^-/-^* neurons

Figure S2. Confirmation of stop cassette deletion in acR3M iMEFs

Figure S3. RNA sequencing analysis of siRNA treated acR3M iMEFs

Figure S4. RNA sequencing analysis of siRNA treated, ZIKV-infected WT neurons

## Acknowledgments

We thank Dr. Daniel Beiting for creating the open-source RNA sequencing learning course, DIY.trancriptomics and the Ram Savan laboratory for generously sharing equipment, reagents and scientific input. S.B.K. also thanks all laboratory members for continued support.

## Author contributions

SBK and AO conceptualized the study and designed experiments. SBK performed most of the experiments and analyzed all the RNA sequencing data. LHC designed and produced all AAV plasmid constructs. JMA advised on RNA sequencing analysis. SDC performed IF staining and confocal imaging and associated quantification. KL technically assisted with cell culture and incucyte assays. BPD provided preliminary data for this project. SBK and AO wrote the manuscript.

## Data and materials availability

*Ripk3^-/-^*mice are available from V.Dixit under a material transfer agreement (MTA) with Genentech.

