## Supplemental Figures for "RIPK3 coordinates RHIM domain-dependent inflammatory transcription in neurons"

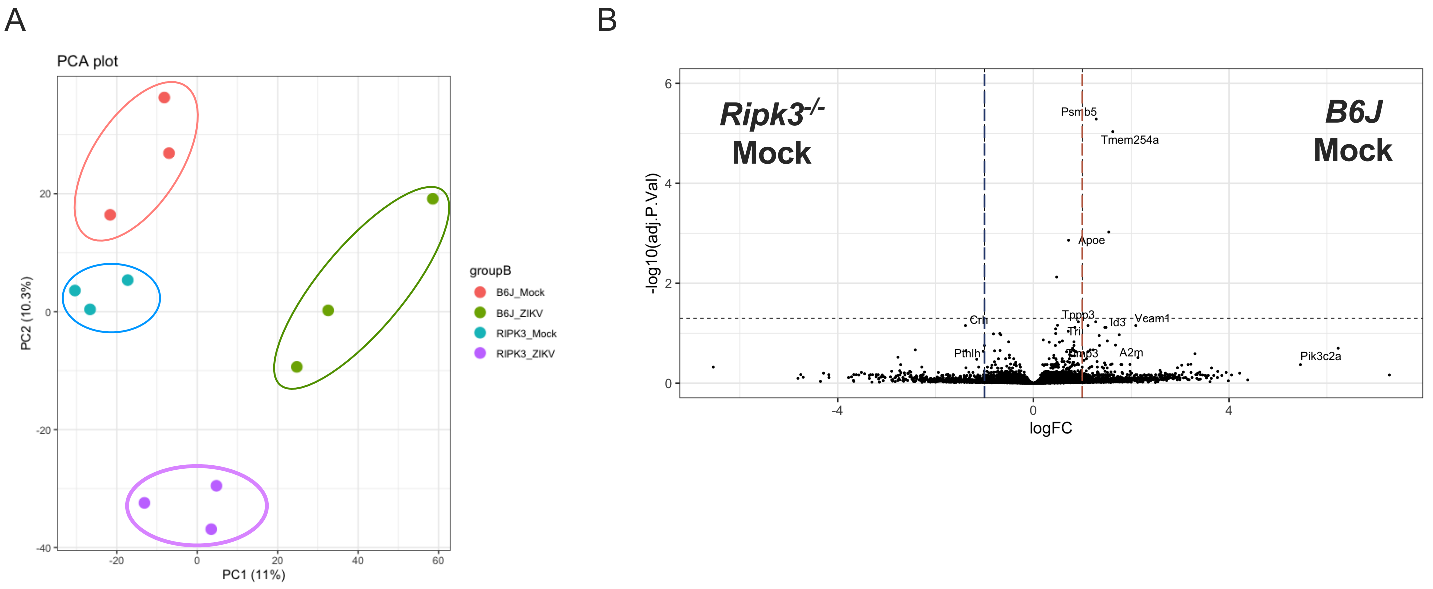


**Supplemental Figure 1: RNA sequencing analysis of ZIKV-infected WT and *Ripk3$^{-/-}$* neurons**

**(A)** Principal component analysis following RNA sequencing of WT (B6J) and *Ripk3$^{-/-}$* neurons 24 hours after ZIKV-MR766 (MOI 0.1) or mock infection (*N*=3 biological replicates). **(B)** Volcano plot depicting differentially expressed genes in WT mock infected samples as compared to *Ripk3$^{-/-}$* mock neuronal samples at 24 hours (*N*=3 biological replicates).

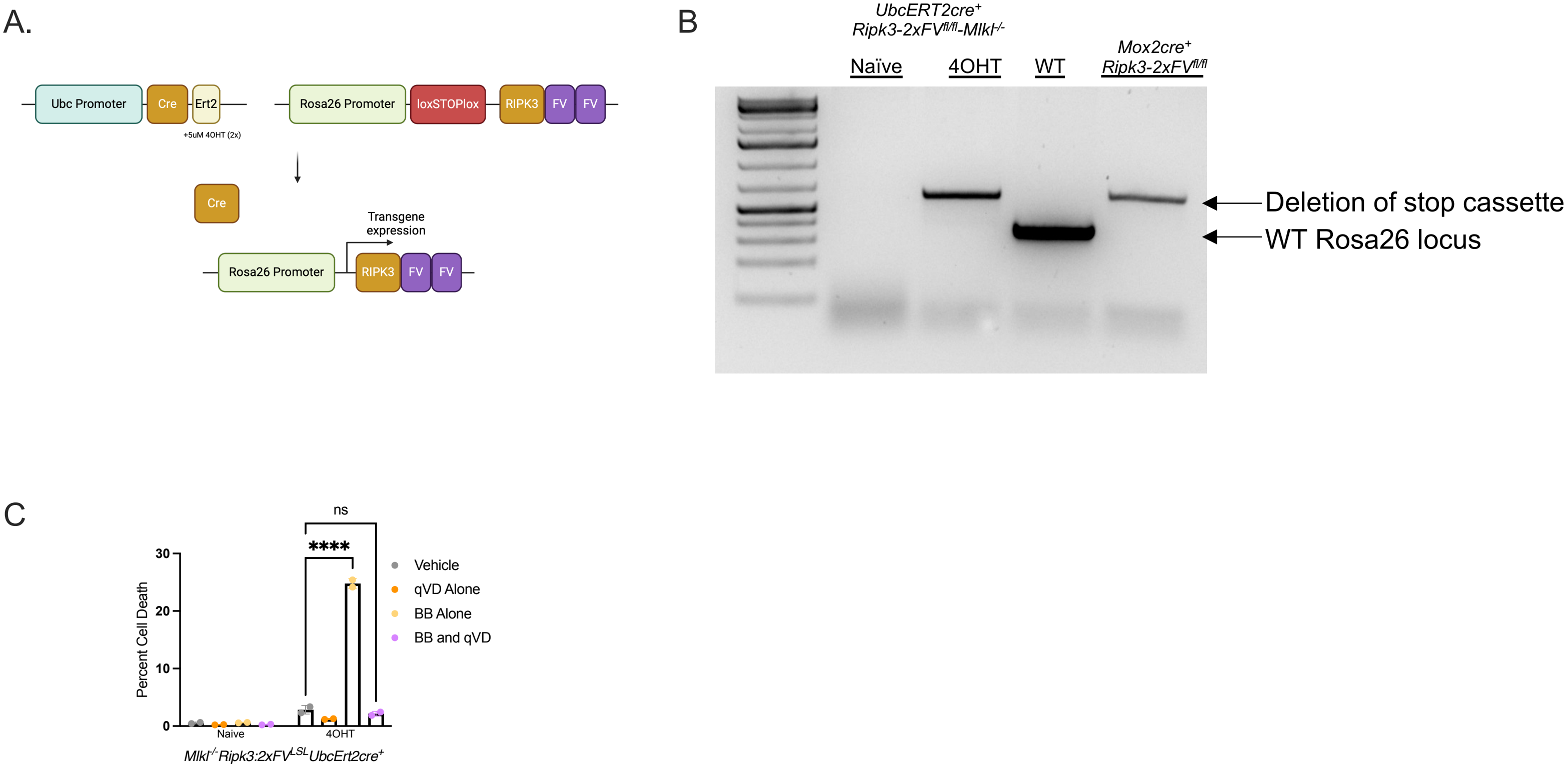


**Supplemental Figure 2: Confirmation of stop cassette deletion in acR3M iMEFs**

**(A)** Schematic for tamoxifen-inducble RIPK3 activatable system in MEFs. **(B)** Successful recombination and confirmation of stop cassette deletion was accomplished by PCR amplification of the Rosa26 locus in acR3M MEFs. WT and *Mox2cre*$^{+}$*Ripk3-2xFV*$^{LSL}$ mouse ear snip lysates were used as a negative and positive control, respectively. **(C)** Percent cell death in acR3M iMEFs 24 hours after treatment with B/B, qVD or B/B and qVD (*N*=2, representative of 3 separate experiments). not significant. ****p<0.0001. Error bars represent SD.

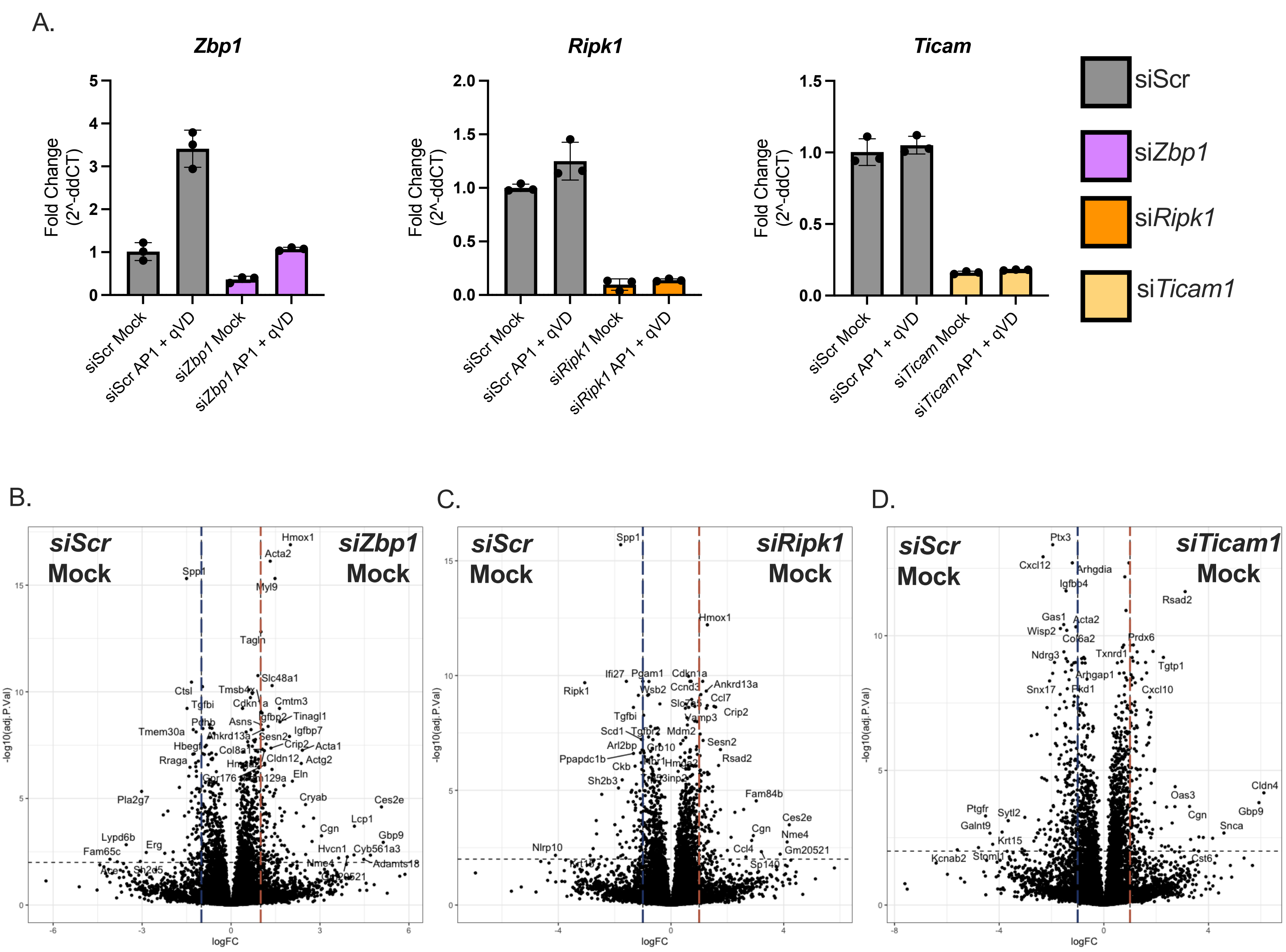


**Supplemental Figure 3: RNA sequencing analysis of siRNA treated acR3M iMEFs**

**(A)** siRNA efficacy was assessed by qRT-PCR in acR3M iMEFs. All samples are normalized to siScr mock values (*N*=3 biological replicates). **(B)** Volcano plot depicting differentially expressed genes in si*Scr* mock infected samples as compared to si*Zbp1* mock acR3M iMEF samples at 24 hours (*N*=3 biological replicates). **(C)** Volcano plot depicting differentially expressed genes in si*Ripk1* mock infected samples as compared to si*Scr* mock acR3M iMEF samples at 24 hours (*N*=3 biological replicates). **(D)** Volcano plot depicting differentially expressed genes in si*Ticam1* mock infected samples as compared to si*Scr* mock acR3M iMEF samples at 24 hours (*N*=3 biological replicates).

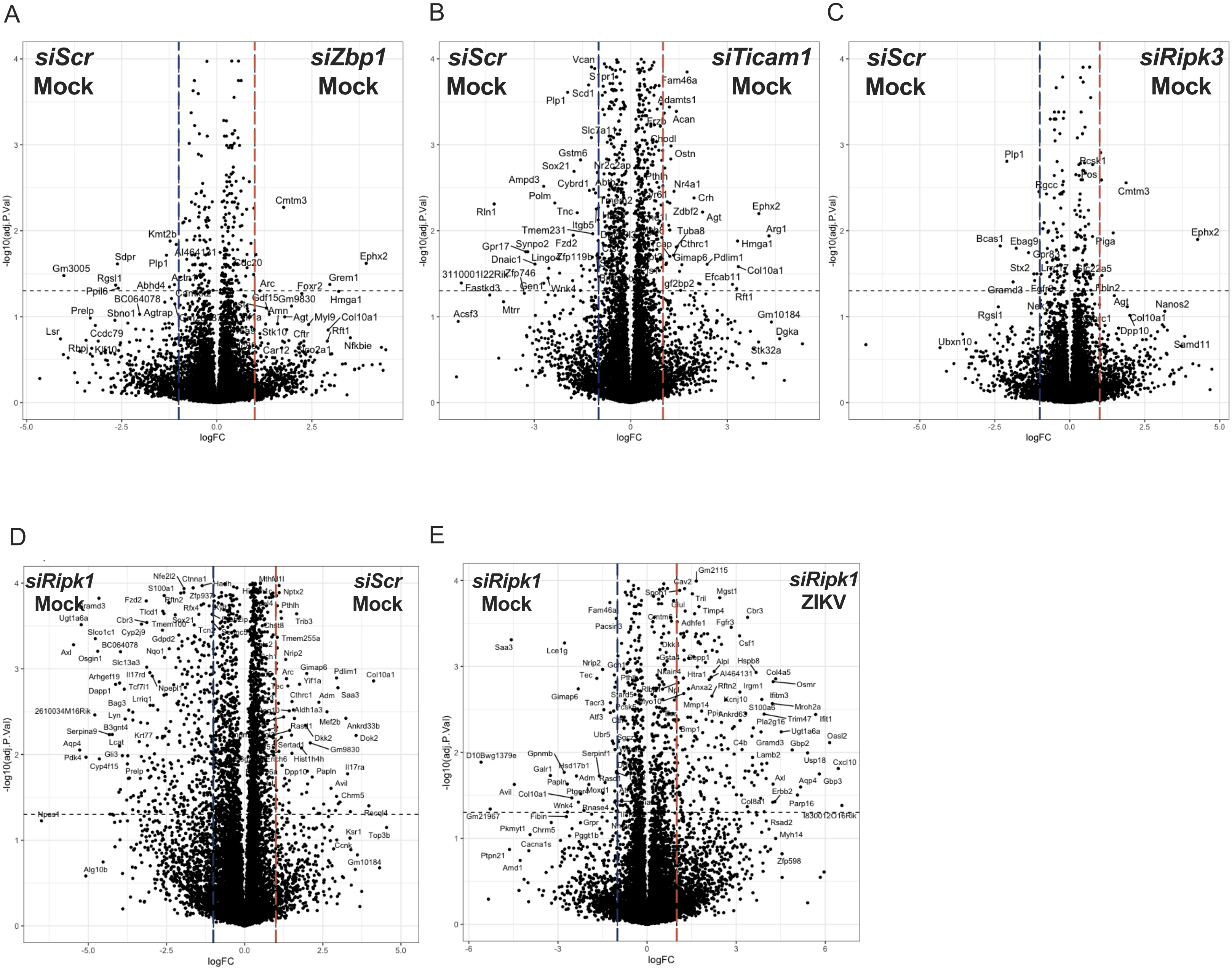


**Supplemental Figure 4: RNA sequencing analysis of siRNA treated, ZIKV-infected WT neurons**

**(A)** Volcano plot depicting differentially expressed genes in si*Zbp1* mock infected samples as compared to si*Scr* mock neuronal samples at 24 hours (*N*=4 biological replicates). **(B)** Volcano plot depicting differentially expressed genes in si*Ticam1* mock infected samples as compared to si*Scr* mock neuronal samples at 24 hours (*N*=4 biological replicates). **(C)** Volcano plot depicting differentially expressed genes in si*Ripk1* mock infected samples as compared to si*Scr* mock neuronal samples at 24 hours (*N*=4 biological replicates). **(D)** Volcano plot depicting differentially expressed genes in si*Scr* mock infected samples as compared to sRipk*1* mock neuronal samples at 24 hours (*N*=4 biological replicates). **(E)** Volcano plot depicting differentially expressed genes in si*Ripk1* ZIKV infected samples as compared to si*Ripk1* mock neuronal samples at 24 hours (*N*=4 biological replicates).
